# Reciprocal allostery arising from a bi-enzyme assembly controls aromatic amino acid biosynthesis in *Prevotella nigrescens*

**DOI:** 10.1101/2021.03.29.437529

**Authors:** Yu Bai, Emily J. Parker

## Abstract

Modular protein assembly has been widely reported as a mechanism for constructing allosteric machinery. Recently, a distinctive allosteric system has been identified in a bi-enzyme assembly comprising a 3-deoxy-d-*arabino* heptulosonate-7-phosphate synthase (DAH7PS) and chorismate mutase (CM). These enzymes catalyze the first and branch point reactions of aromatic amino acid biosynthesis in the bacterium *Prevotella nigrescens* (*Pni*DAH7PS), respectively. The interactions between these two distinct catalytic domains support functional inter-reliance within this bifunctional enzyme. The binding of prephenate, the product of CM-catalyzed reaction, to the CM domain is associated with a striking rearrangement of overall protein conformation that alters the interdomain interactions and allosterically inhibits the DAH7PS activity. In this study, we observed allosteric activation of CM activity in the presence of all DAH7PS substrates. Using small angle X-ray scattering (SAXS) experiments we show that changes in overall protein conformations and dynamics are associated with the presence of different DAH7PS substrates and the allosteric inhibitor prephenate. Furthermore, we have identified an extended interhelix loop located in CM domain, loop_C320-F333_, as a crucial segment for the interdomain structural and catalytic communications. Our results suggest that the dual function enzyme *Pni*DAH7PS contains a reciprocal allosteric system between the two enzymatic moieties, as a result of this bidirectional interdomain communication. This arrangement allows for a complex feedback and feedforward system for control of pathway flux by connecting the initiation and branch point of aromatic amino acid biosynthesis.

---

Allostery is generally defined as the modulation of protein function that arises from ligand binding at a remote site (1,2). Multidomain protein assemblies have been demonstrated as an effective means of delivering allosteric functionality (3-5). Allostery plays an important role in many biosynthetic pathways, where key enzymes are allosterically controlled by pathway end products or intermediates. In some cases, key regulatory enzymes have recruited extra structural elements to the core catalytic unit to acquire allosteric control in response to metabolite concentrations (6-8). Generally, the recruited regulatory elements accommodate effector binding sites and initiate the allosteric signal transduction, delivering activity modulation at the active site (9-13).

Modular assemblies for the delivery of allostery have been well studied in 3-deoxy-d-*arabino*-heptulosonate 7-phosphate synthase (DAH7PS). DAH7PS initiates the shikimate pathway for the biosynthesis of aromatic amino acids (Phe, Tyr, and Trp), by catalyzing the condensation reaction between phosphoenolpyruvate (PEP) and erythrose 4-phosphate (E4P) to produce DAH7P. Allosteric regulation of this gateway enzyme tunes the output of the shikimate pathway to cellular requirements (6-8). DAH7PS typically exists as a homotetramer or homodimer of catalytic subunits that comprise a (β/α)_8_ TIM barrel (11,14-18). The acquisition of allostery by the DAH7PS enzyme family involves diverse additions to the core barrel (19), amongst which the adornments of discrete structural modules to either end of the DAH7PS domain are observed in one subtype of DAH7PS enzymes (type Iβ). The allosteric response conferred by these allosteric modules is characterized by dramatic conformational changes in response to allosteric ligand binding (11,16). In addition to protein modules that solely deliver allostery, type Iβ DAH7PS proteins are also observed to be covalently linked, due to gene fusion, to another enzyme in aromatic amino acid biosynthesis – chorismate mutase (CM) (10,11,20).

CM is a pivotal enzyme at a branch point of the shikimate pathway, catalyzing the transformation of chorismate to prephenate, which ultimately delivers Phe and Tyr. Two main classes of CM enzymes have been identified (21,22). The CM enzymes associated with DAH7PS belong to the AroQ subtype and exert function as a homodimer of three-helix bundles (23). The AroQ CM is fused to either the N-terminus or C-terminus of DAH7PS. For the N-terminal CM-linked DAH7PS enzymes, such as the DAH7PS from *Bacillus subtilis* (*Bsu*DAH7PS), *Listeria monocytogenes* (*Lmo*DAH7PS), and *Geobacillus sp*. (*Gsp*DAH7PS), the role of CM as an allosteric domain is well demonstrated (10,11,20). The crystal structure of *Gsp*DAH7PS shows that the N-terminal CM of each DAH7PS barrel pairs with its diagonally opposed counterpart and form a dimer at either side of the DAH7PS tetrameric core (Figure 1A) (11). The binding of prephenate (or chorismate) to the active site of CM is associated with a conformational rearrangement where the CM dimers shift towards a tighter association with the tetrameric DAH7PS core. This significant conformational change results in the occlusion of the entrances of DAH7PS active sites and disrupts catalysis (Figure 1A) (11). In this complex system, although the CM is catalytically functional, it appears to work as nothing more than a regulatory module to provide a gating mechanism and the CM active site acts as an allosteric effector binding site. In addition to the AroQ CM, a monofunctional unregulated CM belonging to AroH subtype is encoded in the organisms with N-terminal CM-linked DAH7PS. This AroH CM exhibits a higher enzymatic activity and is predicted to be the main *in vivo* catalyst for the production of prephenate (24).

**Figure 1:**
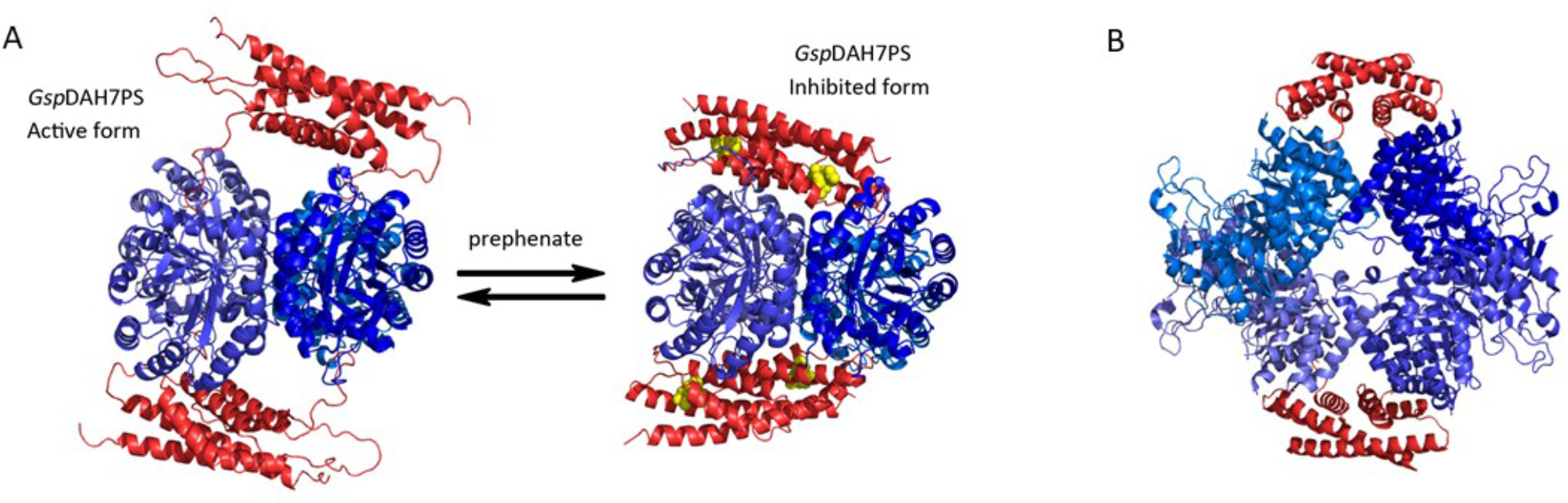
Crystal structures and homology model illustrating the allosteric regulations of *Gsp*DAH7PS and the architecture of *Mtu*DAH7PS-CM complex. DAH7PS barrels are shown in blue, and CM domains are colored in red. **(A)** The binding of prephenate (yellow spheres) shifts the conformational population of *Gsp*DAH7PS resulting in inhibition of DAH7PS activity (11). **(B)** Non-covalent complex between *Mtu*DAH7PS and the intracellular *Mtu*CM (30).

In contrast, the C-terminal CM-linked DAH7PS is often found as the only source of CM activity in the organisms where it is identified from according to classification of CM enzymes retrieved from Pfam database (25). Compared with other well-studied DAH7PS proteins, these enzymes adopt distinctive machinery for the delivery of both allostery and catalysis, as reported by our recent study on a DAH7PS from *P. nigrescens* (*Pni*DAH7PS) (26). *Pni*DAH7PS is a bifunctional enzyme consisting of an N-terminal DAH7PS and a C-terminal CM domain, and separation of the two domains results in a dramatic attenuation of both enzymatic activities. The structural investigation into this protein revealed that it displays a unique homodimeric assembly with dimerization of the helical CM domains, and no direct contact between DAH7PS barrels (26). Moreover, DAH7PS activity is allosterically inhibited by prephenate (26). This allosteric inhibition is accompanied by a striking conformational change that alters the interdomain interactions between DAH7PS and CM (26), which indicates that a manipulation of the interdomain interaction is the mechanism underpinning allosteric inhibition.

A similar functional reliance between the DAH7PS and CM is observed in the noncovalent DAH7PS-CM complex found in *Corynebacterium glutamicum* (*Cgl*DAH7PS-CM) (27) and *Mycobacterium tuberculosis* (*Mtu*DAH7PS-CM) (28). In the *Mtu*DAH7PS-CM complex, CM is activated upon complexation with DAH7PS and inhibited by the binding of allosteric inhibitors, Tyr and Phe, at allosteric binding sites within each DAH7PS subunit (29-31). The structure of the *Mtu*DAH7PS-CM complex showed that the DAH7PS and CM subunits interact with each other extensively and form a stable interface (Figure 1B) (28,30). Molecular dynamics simulations suggest that the interface between DAH7PS and CM subunits is noticeably more flexible when the inhibitors are bound, which may result in a less stable DAH7PS-CM interaction (32), suggesting a crucial role of this stable interface for delivering CM function.

In contrast to the *Mtu*DAH7PS-CM complex, the functional enhancements that arise from association are mutual between DAH7PS and CM activities of *Pni*DAH7PS, which suggests an extensive DAH7PS-CM interaction and communication pathway between the two functional domains. Furthermore, SAXS data of ligand-free *Pni*DAH7PS showed that it is highly flexible and able to adopt a broad range of relative domain conformations (26), indicating a lack of stable domain interface, unlike *Mtu*DAH7PS-CM. How then does such a flexible structure of *Pni*DAH7PS deliver the functional communication between two active sites? To address this question, we conducted a series of biochemical and structural investigations on *Pni*DAH7PS in the presence of different ligands. Our studies reveal dynamic and variable interdomain interactions for *Pni*DAH7PS are associated with the DAH7PS-catalyzed reaction and that these result in reciprocal allosteric regulation between the DAH7PS and CM active sites.

## RESULTS

### DAH7PS allosterically activates CM

Previous kinetic studies revealed the interdependency of the DAH7PS and CM catalytic functions of *Pni*DAH7PS and demonstrated that prephenate binding to the CM domain allosterically inhibits DAH7PS activity (26). To understand the domain interdependency and the full allosteric functionality of this protein, we investigated the effect of substrate binding and catalysis in the DAH7PS active site on the catalysis of the CM reaction. The catalytic rate for the CM reaction was measured directly in the presence of the DAH7PS substrates and metal cofactor, either individually or in combination (Figure 2). Interestingly, we observed a greater than two-fold increase in the rate of chorismate consumption in the presence of all DAH7PS reaction components (PEP/Mn^2+^/E4P, Figure 2A). It should be noted that under this experimental condition, catalysis was occurring simultaneously at both CM and DAH7PS active sites. This observation implies that there is an allosteric activation effect, exerted by the occupancy and catalysis at the DAH7PS active site, on the CM catalytic function for *Pni*DAH7PS.

**Figure 2:**
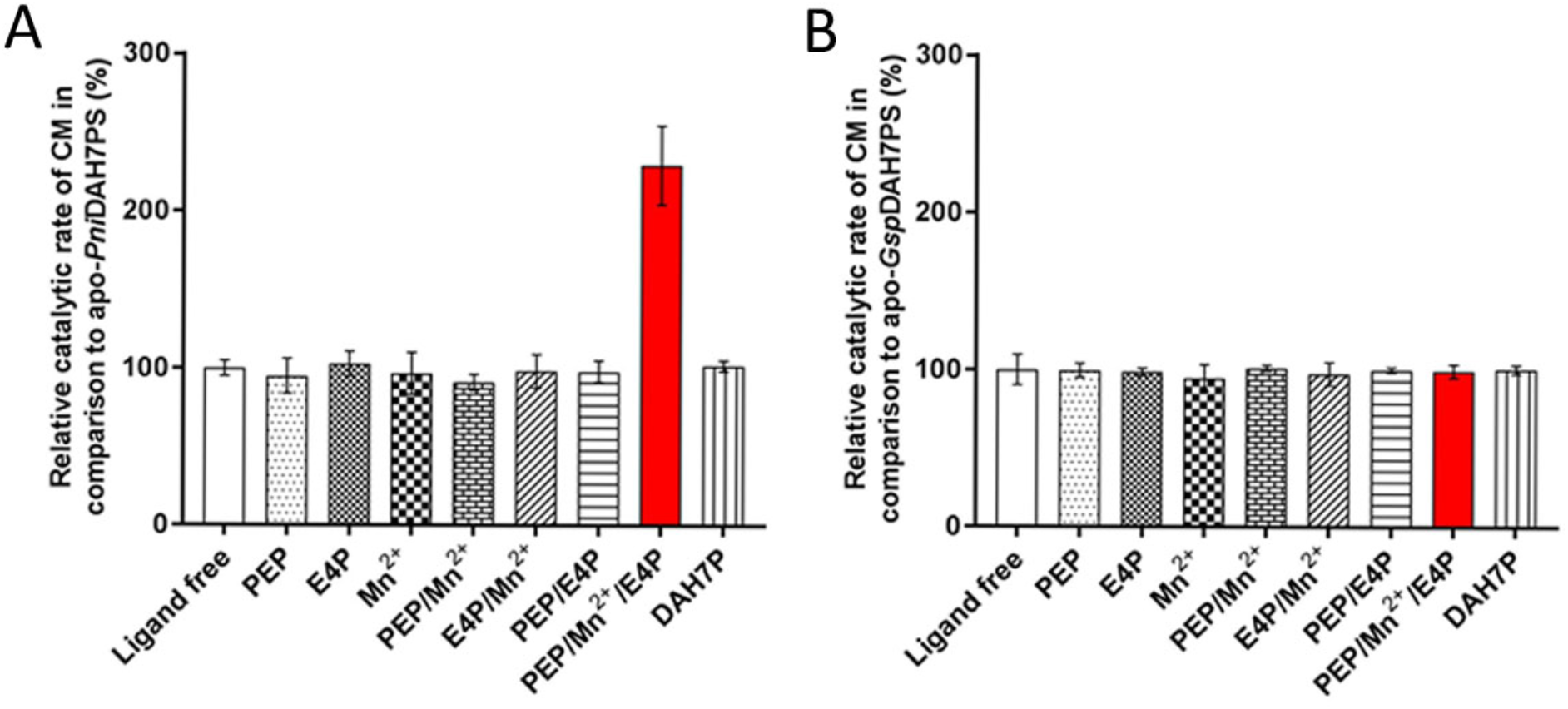
The presence of PEP, Mn^2+^ and E4P boosts the CM catalytic rate of *Pni*DAH7PS. Error bars represent the standard deviation of triplicate measurements. The CM catalytic rates of *Pni*DAH7PS **(A)** and *Gsp*DAH7PS **(B)** were examined respectively in the presence of a variety of combinations of DAH7PS reaction substrates, product and Mn^2+^ ion as indicated. Direct interference by the components of DAH7PS reaction in the detection of CM activity, was excluded, as detailed in the methods section.

In contrast, when the same analysis was carried out for the N-terminal CM-linked *Gsp*DAH7PS, no variation in the CM reaction rate was detected with any combination of the DAH7PS substrates (Figure 2B). This finding indicates that the observed allosteric activation of CM activity by binding and/or catalytic events at the DAH7PS reaction is specific to *Pni*DAH7PS.

### Conformational changes of PniDAH7PS are related to the occupancy of the DAH7PS active site

The structural effects of DAH7PS substrates on *Pni*DAH7PS were inspected using SAXS. These measurements were achieved by passing the enzyme through a size exclusion column, using an elution buffer containing combinations of PEP, E4P, and MnSO_4_, prior to exposure to the X-ray beam. The scattering profile from this protein is denoted here as *Pni*DAH7PS_rxn_. With all three reaction components present, the protein displays a more compact structure, compared to that of the apo-enzyme, as evidenced by the reduced *R*_*g*_, *D*_max_ and *V*_p_ values (37.7 ± 0.6 Å, 140.2 Å and 121 nm^3^ respectively, Table 1). Furthermore, the Kratky and Porod-Debye plots derived from the scattering data indicates a decreased flexibility in the protein (Figure 3C). It should be noted that, given that the DAH7PS-catalyzed reaction is underway in the presence of both substrates and metal ion (PEP, E4P and Mn^2+^), the SAXS scattering of *Pni*DAH7PS_rxn_ arises from the average conformation of the protein through the catalytic cycle.

**Table 1:** SAXS parameters of apo-*Pni*DAH7PS, *Pni*DAH7PS_PM_ and *Pni*DAH7PS_rxn_.

|  | Ligands present |  |  |
| --- | --- | --- | --- |
|  | Ligand-free | PEP/Mn<sup>2+</sup> | PEP/Mn<sup>2+</sup>/E4P |
| **Guinier Analysis** |  |  |  |
| $R_g$ (Å) | 42.0 ± 1.2 | 39.4 ± 0.8 | 35.8 ± 1.0 |
| $I(0)$ (cm<sup>-1</sup>) | 0.08 ± 0.00 | 0.07 ± 0.00 | 0.07 ± 0.00 |
| Correlation coefficient, $R^2$ | 0.99 | 0.99 | 0.99 |
| **Pair Wise Distribution Analysis** |  |  |  |
| $R_g$ (Å) | 44.1 ± 0.2 | 41.0 ± 0.1 | 37.7 ± 0.6 |
| $I(0)$ (cm<sup>-1</sup>) | 0.08 ± 0.00 | 0.07 ± 0.00 | 0.07 ± 0.00 |
| $D_{max}$ (Å) | 152.3 | 142.7 | 140.2 |
| $V_p$ (nm<sup>3</sup>) | 137 | 139 | 121 |
| **$MM_{porod}$** |  |  |  |
| $MM$ (kDa) | 81 | 82 | 71 |
| Number of Subunits | 2 | 2 | 2 |

$V_p$ : Porod volume; $MM_{porod}$ : molecular mass estimated from Porod volume. The uncertainties represent standard deviations estimated by the error propagation from the experimental data.

**Figure 3:**
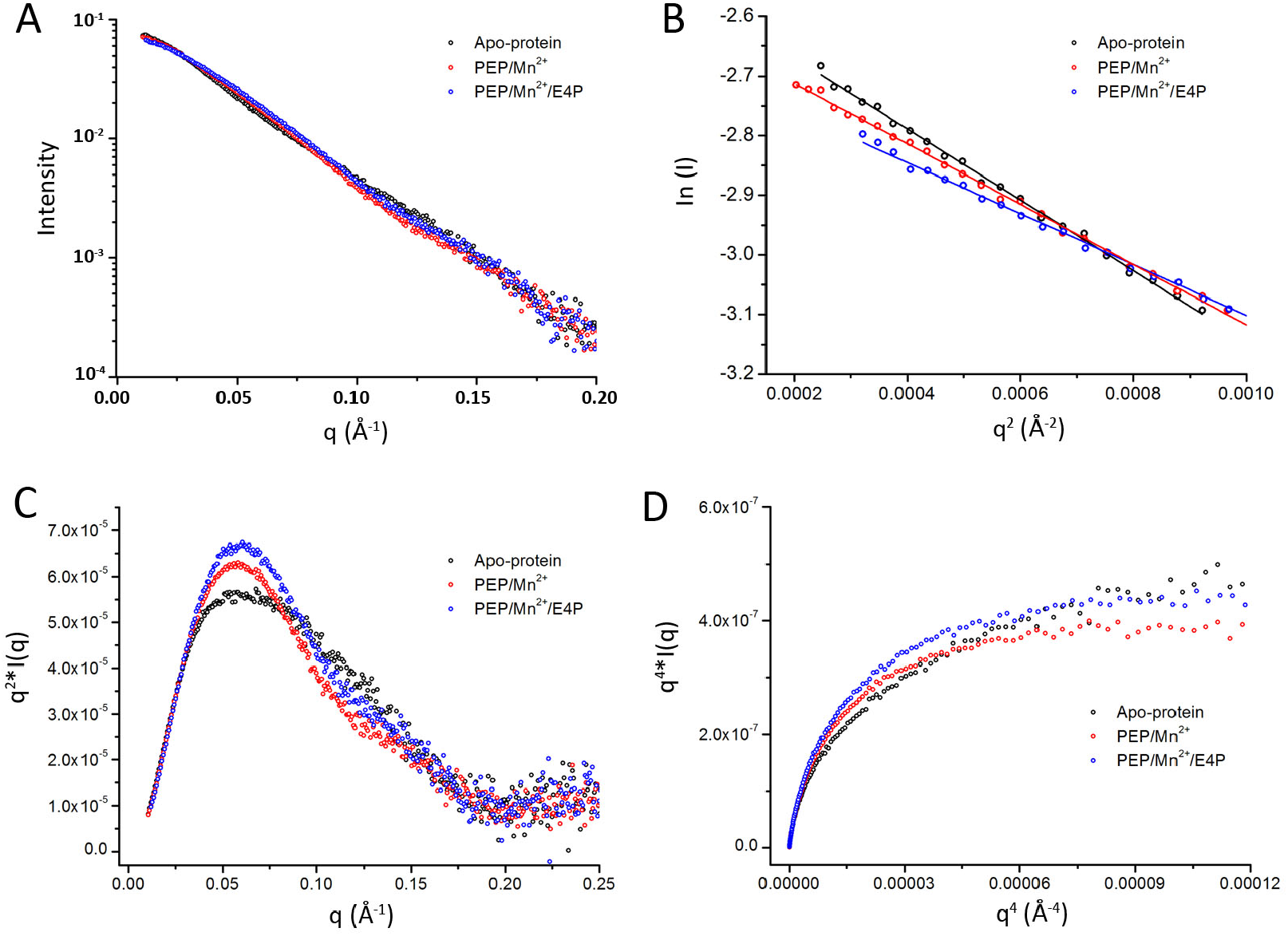
Analysis of the SAXS data of apo-*Pni*DAH7PS (black) and *Pni*DAH7PS_PM_ (red) and *Pni*DAH7PS_rxn_ (blue). **(A)** SAXS profiles (log *I*(*q*) versus *q*). **(B)** Guinier plots (ln *I*(*q*) versus *q*^2^). **(C)** Kratky plot (*q*^2^\**I*(*q*) versus *q*). **(D)** Porod-Debye plot (*q*^4^\**I*(*q*) versus *q*^4^) limited to the range of the SAXS data for which the Guinier linearity was observed.

Further SAXS analysis revealed that the addition of both PEP and Mn^2+^ also alters the scattering from the protein, implying that this liganded *Pni*DAH7PS (*Pni*DAH7PS_PM_) adopts a more compact structure than the apo-protein, albeit not as compressed as that of the protein in the presence of all DAH7PS reaction components (Table 1 and Figure 3B). In addition to reductions of *R*_g_ and *D*_max_ values, structural rigidification was also observed, as indicated by the Kratky and Porod-Debye transformations (Figure 3C, D).

To visualize the conformational changes indicated by the scattering profiles described above, rigid body modelling was carried out using models for *Pni*DAH7PS^D^ and *Pni*DAH7PS^CM^ (26) and the real space information extracted from the SAXS data. Rigid body models for apo-*Pni*DAH7PS, *Pni*DAH7PS_PM_ and *Pni*DAH7PS_rxn_ yield reasonable fits (χ^2^ = 0.3, 1.47 and 1.43 respectively) to their corresponding experimental SAXS data (Figure 4). Both the models of *Pni*DAH7PS_PM_ and *Pni*DAH7PS_rxn_ adopt more compact spatial organizations compared to that of apo-*Pni*DAH7PS (Figure 4). It is of particular note that in the model for *Pni*DAH7PS_rxn_ the two DAH7PS barrels are predicted to be positioned in close proximity to the CM dimer, as demonstrated by the decreased distance, from 39 Å to 27 Å, between residues R165 (NH1 atom) and R312 (NH1 atom) in the same chain which are respectively situated at the centers of DAH7PS and CM active sites. This model might provide a structural rationale for forming an extensive interdomain interface and subsequently promoting both CM and DAH7PS activities under these conditions (Figure 4C). These SAXS-based observations and analyzes suggest that a diverse range of interdomain interactions are possible and that conformational plasticity is associated with the DAH7PS catalytic cycle.

**Figure 4:**
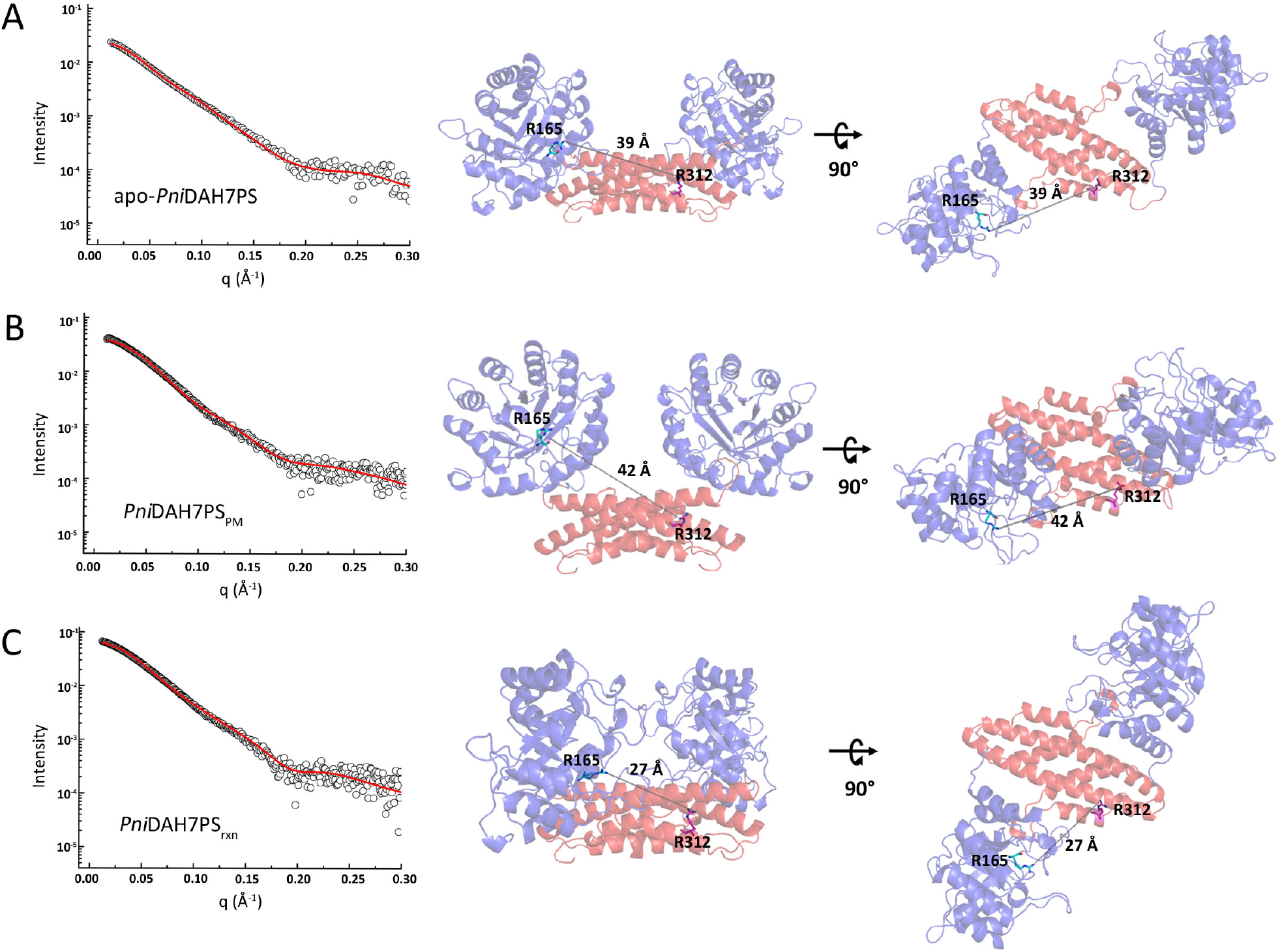
Rigid body models for apo-*Pni*DAH7PS, *Pni*DAH7PS_PM_ and *Pni*DAH7PS_rxn_ derived SAXS data. DAH7PS and CM domains are respectively colored blue and red, and the distances between the NH1 atom of R165 (cyan stick) and the NH1 atom of R312 (pink stick) are indicated as grey dash lines. **(A)** Rigid body model of apo-*Pni*DAH7PS and the fit to the experimental SAXS data (*χ*^2^= 0.3). **(B)** Rigid body model of *Pni*DAH7PS_PM_ and the fit to the experimental SAXS data (*χ*^2^=1.47). **(C)** Rigid body model of *Pni*DAH7PS_rxn_ and the fit to the experimental SAXS data (*χ*^2^=1.43).

### A loop truncation in the CM domain interferes with catalysis and conformation of PniDAH7PS

A multisequence alignment (MSA) of a number of AroQ CM enzymes, including the independent monofunctional CM enzymes from *Pyrococcus furiosus* (*Pfu*CM) and *Aeropyrum pernix* (*Ape*CM), and the CM domains of *Pni*DAH7PS and the C-terminal CM-linked DAH7PS from *Porphyromonas gingivalis, Pgi*DAH7PS (*Pni*DAH7PS^CM^ and *Pgi*DAH7PS^CM^), was constructed. This MSA and structural comparison revealed that in comparison to the discrete AroQ CM proteins, *Pni*DAH7PS^CM^ contains a ∼10 residue extension to the α2-α3 loop, loop_C320-F333_ (Figure 5). This loop extension is also observed in other CM domains linked to the C-terminal of a DAH7PS catalytic barrel.

**Figure 5:**
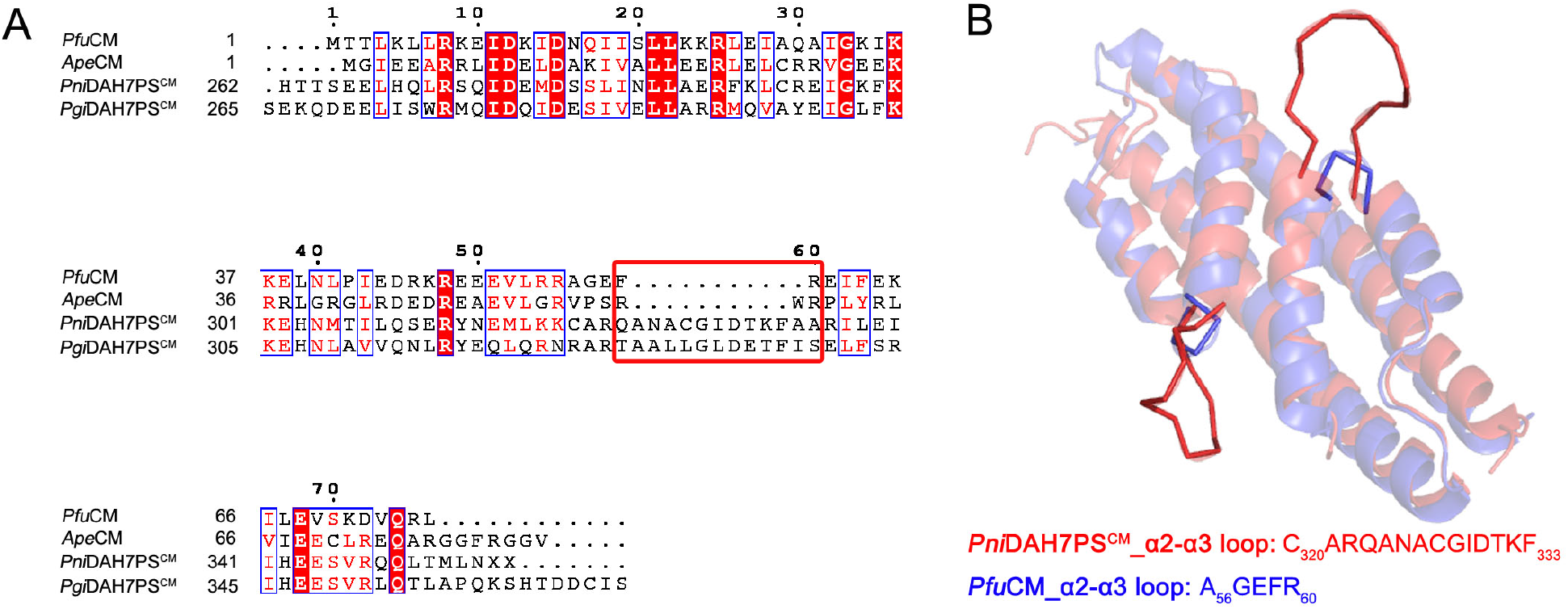
Sequence and structural alignments predict a loop extension within *Pni*DAH7PS^CM^. **(A)** The sequence alignment of *Pni*DAH7PS^CM^ with *Pgi*DAH7PS^CM^ with monofunctional AroQ CMs. The red rectangle highlights the unaligned region that is predicted to be an extended α2-α3 loop. **(B)** The structural alignment between *Pfu*CM (crystal structure, shown in blue cartoon) and *Pni*DAH7PS^CM^ (homology model, shown in red cartoon), showing the extended loop between helices 2 and 3.

To determine whether this extended loop affects the function and structure of *Pni*DAH7PS, a variant of *Pni*DAH7PS with a shortened loop (*Pni*DAH7PS^Δ5AA^) was generated, by removing five residues (CGIDT) from the loop. Kinetic measurements of this variant demonstrated a considerable loss in the CM catalytic efficiency, as evidenced by the decrease in the *k*_cat_/*K*_M_ value for chorismate from 0.22 to 5.7 × 10^−2^ μM^-1^* s^-1^ (Table 2). Notably, in addition to impaired CM catalysis, the DAH7PS activity of *Pni*DAH7PS^Δ5AA^ was also significantly attenuated; the apparent catalytic efficiency, *k*_cat_/*K*_m_ values, for PEP and E4P were decreased respectively from 0.41 and 0.29 μM^-1^* s^-1^ to 4.7 × 10^−2^ and 1.8 × 10^−2^ μM^-1^* s^-1^ (Table 2).

**Table 2:** Kinetic characteristics of the DAH7PS and CM functions of *Pni*DAH7PS^Δ5AA^, and the kinetic parameters of *Pni*DAH7PS, *Pni*DAH7PS^D^, and *Pni*DAH7PS^CM^ for comparison.

| Enzyme | DAH7PS activity |  |  |  |  | CM activity |  |  |
| --- | --- | --- | --- | --- | --- | --- | --- | --- |
| | $K_M^{\text{PEP}}$ | $K_M^{\text{E4P}}$ | $k_{\text{cat}}$ | $k_{\text{cat}}/K_M^{\text{PEP}}$ | $k_{\text{cat}}/K_M^{\text{E4P}}$ | $K_M$ | $k_{\text{cat}}$ | $k_{\text{cat}}/K_M$ |
| | ($\mu\text{M}$) | ($\mu\text{M}$) | ($\text{s}^{-1}$) | ($\mu\text{M}^{-1} \cdot \text{s}^{-1}$) | ($\mu\text{M}^{-1} \cdot \text{s}^{-1}$) | ($\mu\text{M}$) | ($\text{s}^{-1}$) | ($\mu\text{M}^{-1} \cdot \text{s}^{-1}$) |
| *Pni*DAH7PS (26) | 41 ± 4 | 58 ± 6 | 16.8 ± 0.5 | 0.41 | 0.29 | 7.6 ± 0.6 | 1.68 ± 0.06 | 0.22 |
| *Pni*DAH7PS<sup>Δ5AA</sup> | 212 ± 51 | 556 ± 77 | 11 ± 1 | 4.7 × 10<sup>-2</sup> | 1.8 × 10<sup>-2</sup> | 21 ± 2 | 1.20 ± 0.02 | 5.7 × 10<sup>-2</sup> |
| *Pni*DAH7PS<sup>D</sup> (26) | 262 ± 25 | 652 ± 60 | 1.6 ± 0.1 | 6.1 × 10<sup>-3</sup> | 2.5 × 10<sup>-3</sup> | NA | NA | NA |
| *Pni*DAH7PS<sup>CM</sup> (26) | NA | NA | NA | NA | NA | 84 ± 5 | 2.01 ± 0.07 | 2.4 × 10<sup>-2</sup> |

The Michaelis constants for the DAH7PS activity were obtained via fitting kinetic data to the equation of Alberty (as detailed in the method section). The uncertainty values represent the standard deviation from triplicate measurements.

In parallel, we examine the structural properties of *Pni*DAH7PS^Δ5AA^ using SAXS experiments. The scattering data reveals that apo-*Pni*DAH7PS^Δ5AA^ retains a flexible and elongated homodimeric structure (Table 3). However, the increased *R*_g_ and *V*_p_ values of apo-*Pni*DAH7PS^Δ5AA^ (Table 3), compared to those for wild-type apo-*Pni*DAH7PS, indicate that the loop truncation increased the average size of the *Pni*DAH7PS homodimer (Figure 6D). Following the addition of PEP and Mn^2+^, the conformation of *Pni*DAH7PS^Δ5AA^ 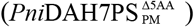 became more compact, presenting a similar average size to that of *Pni*DAH7PS_PM_ (Table 3). Whereas the Kratky and Porod-Debye transformations of 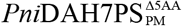 showed considerable flexibility, which differs from the rigidification of the wild-type protein upon the binding of PEP and Mn^2+^ (Figure 6H, I). Furthermore, with the presence of PEP/Mn^2+^/E4P, the conformation of *Pni*DAH7PS^Δ5AA^ 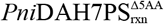 displayed a considerable increase in the protein size compared with that for 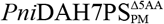 (Table 3). The flexibility of 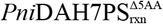 remained high, as demonstrated by the Kratky and Porod-Debye plots (Figure 6K, L). Overall, the truncated variant appears to respond differently from the wild-type enzyme to the presence of either PEP/Mn^2+^ or PEP/Mn^2+^/E4P and maintains high flexibility under all the conditions. These observations imply that the loop truncation likely weakens DAH7PS-CM interdomain interactions, which in turn interferes with the catalytic properties of *Pni*DAH7PS^Δ5AA^.

**Table 3:** SAXS parameters of apo-*Pni*DAH7PS^Δ5AA^, 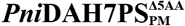 and 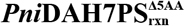.

|  | Ligand present |  |  |
| --- | --- | --- | --- |
|  | Ligand-free | PEP/Mn<sup>2+</sup> | PEP/Mn<sup>2+</sup>/E4P |
| **Guinier Analysis** |  |  |  |
| $R_g$ (Å) | 45.6 ± 0.8 | 39.9 ± 1.0 | 42.3 ± 1.7 |
| $I(0)$ (cm<sup>-1</sup>) | 0.08 ± 0.00 | 0.07 ± 0.00 | 0.08 ± 0.00 |
| Correlation coefficient, R<sup>2</sup> | 0.99 | 0.99 | 0.99 |
| **Pair Wise Distribution Analysis** |  |  |  |
| $R_g$ (Å) | 46.1 ± 0.6 | 41.1 ± 0.5 | 43.4 ± 0.4 |
| $I(0)$ (cm<sup>-1</sup>) | 0.08 ± 0.00 | 0.07 ± 0.00 | 0.08 ± 0.00 |
| $D_{max}$ (Å) | 150.5 | 140.1 | 146.5 |
| $V_p$ (nm<sup>3</sup>) | 175 | 128 | 143 |
| **$MM_{porod}$** |  |  |  |
| MM (kDa) | 103 | 75 | 84 |
| Number of Subunits | 2 | 2 | 2 |

$V_p$ : Porod volume; $MM_{porod}$ : molecular mass estimated from Porod volume. The uncertainties represent standard deviations estimated by the error propagation from the experimental data.

**Figure 6:**
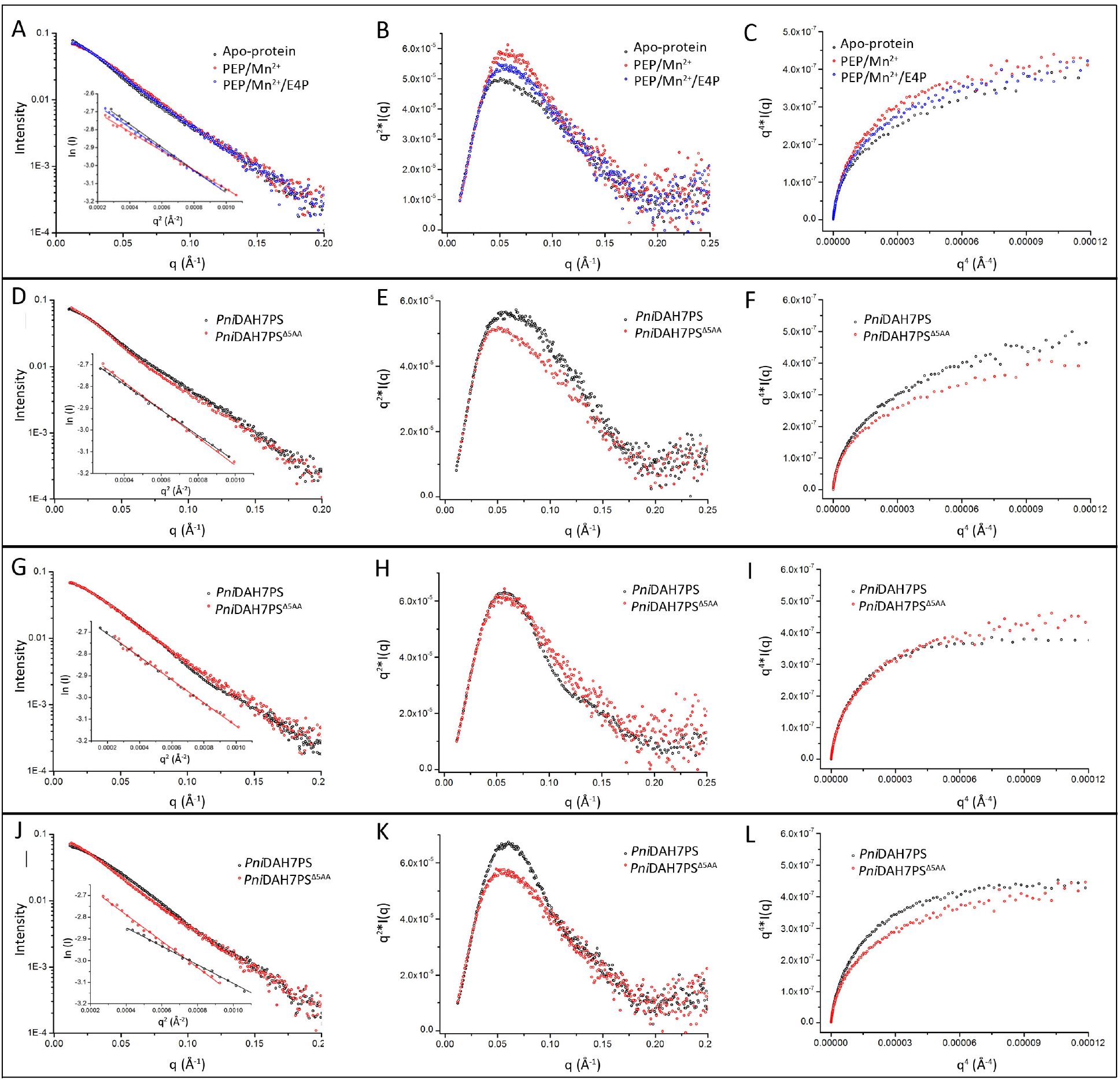
The analysis of the SAXS data for *Pni*DAH7PS^Δ5AA^, and the comparison between the conformational changes of *Pni*DAH7PS^Δ5AA^ and *Pni*DAH7PS resulted from the addition of either PEP/Mn^2+^ or PEP/Mn^2+^/E4P. **The first panel** shows the SAXS profiles **(A)** and Kratky and Porod-Debyem plots **(B and C)** of apo-*Pni*DAH7PS^Δ5AA^ (black), 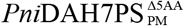 (red) and 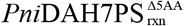 (blue) respectively. **The second panel** compares the SAXS profiles **(D)**, the Kratky and Porod-Debye plots **(E and F)** of apo-*Pni*DAH7PS^Δ5AA^ (red) with that of apo-*Pni*DAH7PS (black). **The third panel** compares the SAXS profiles **(G)**, the Kratky and Porod-Debye plots **(H and I)** of 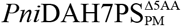 (red) with that of *Pni*DAH7PS_PM_ (black). **The fourth panel** compares the SAXS profiles **(J)**, the Kratky and Porod-Debye plots **(K, L)** of 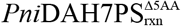 (red) with that of *Pni*DAH7PS_rxn_ (black).

### Prephenate restricts the conformational change of PniDAH7PS and disrupts the catalytic cycle of DAH7PS reaction

Previous studies only demonstrated the inhibitory effect of prephenate on *Pni*DAH7PS (26). Here we conducted kinetic assays for the DAH7PS activities in the presence of prephenate and observed higher *K*_m_ values for both PEP and E4P and a reduced *k*_cat_, which led to the substantial reduction in catalytic efficiency (Table 4).

**Table 4:** Kinetic parameters for the DAH7PS activity of *Pni*DAH7PS in either the absence or the presence of 100 µM prephenate.

| Enzyme | $K_M^{\text{PEP}}$<br>(μM) | $K_M^{\text{E4P}}$<br>(μM) | $k_{\text{cat}}$<br>($\text{s}^{-1}$) | $k_{\text{cat}}/K_M^{\text{PEP}}$<br>($\mu\text{M}^{-1} \cdot \text{s}^{-1}$) | $k_{\text{cat}}/K_M^{\text{E4P}}$<br>($\mu\text{M}^{-1} \cdot \text{s}^{-1}$) |
| --- | --- | --- | --- | --- | --- |
| **Apo-*Pni*DAH7PS (26)** | 41 ± 4 | 58 ± 6 | 16.8 ± 0.5 | 0.41 | 0.29 |
| **Prephenate-*Pni*DAH7PS** | 102 ± 34 | 366 ± 51 | 4.8 ± 0.5 | $4.7 \times 10^{-2}$ | $1.3 \times 10^{-2}$ |

The Michaelis constants for the DAH7PS activity were obtained via fitting kinetic data to the equation of Alberty (as described in the method section). The uncertainty values represent the standard deviation from triplicate measurements.

We previously reported that upon the binding of prephenate, the conformational distribution of apo-*Pni*DAH7PS shifts towards a more compact and rigid average structure (26). This prephenate-bound conformation is more compact than that of *Pni*DAH7PS observed in the presence of either PEP/Mn^2+^ or PEP/Mn^2+^/E4P, with a *R*_g_ value of 33Å (Table 5). To further probe the effect of prephenate on the conformational variations of *Pni*DAH7PS associated with the occupancy of the DAH7PS active site, SAXS data for *Pni*DAH7PS was collected in the presence of both prephenate and combinations of DAH7PS active site ligands. Intriguingly, the distinct conformational populations observed for the prephenate free *Pni*DAH7PS (Table 1) are no longer apparent for the prephenate bound enzymes (Table 5 and Figure 7). When prephenate is present, *Pni*DAH7PS maintains the compact and rigid conformations under different combinations of DAH7PS substrates (Table 5 and Figure 7). This observation implies that prephenate binding not only shifts the conformational distribution of *Pni*DAH7PS but also restricts the redistribution of conformations associated with the occupation of the DAH7PS active site.

**Table 5:**
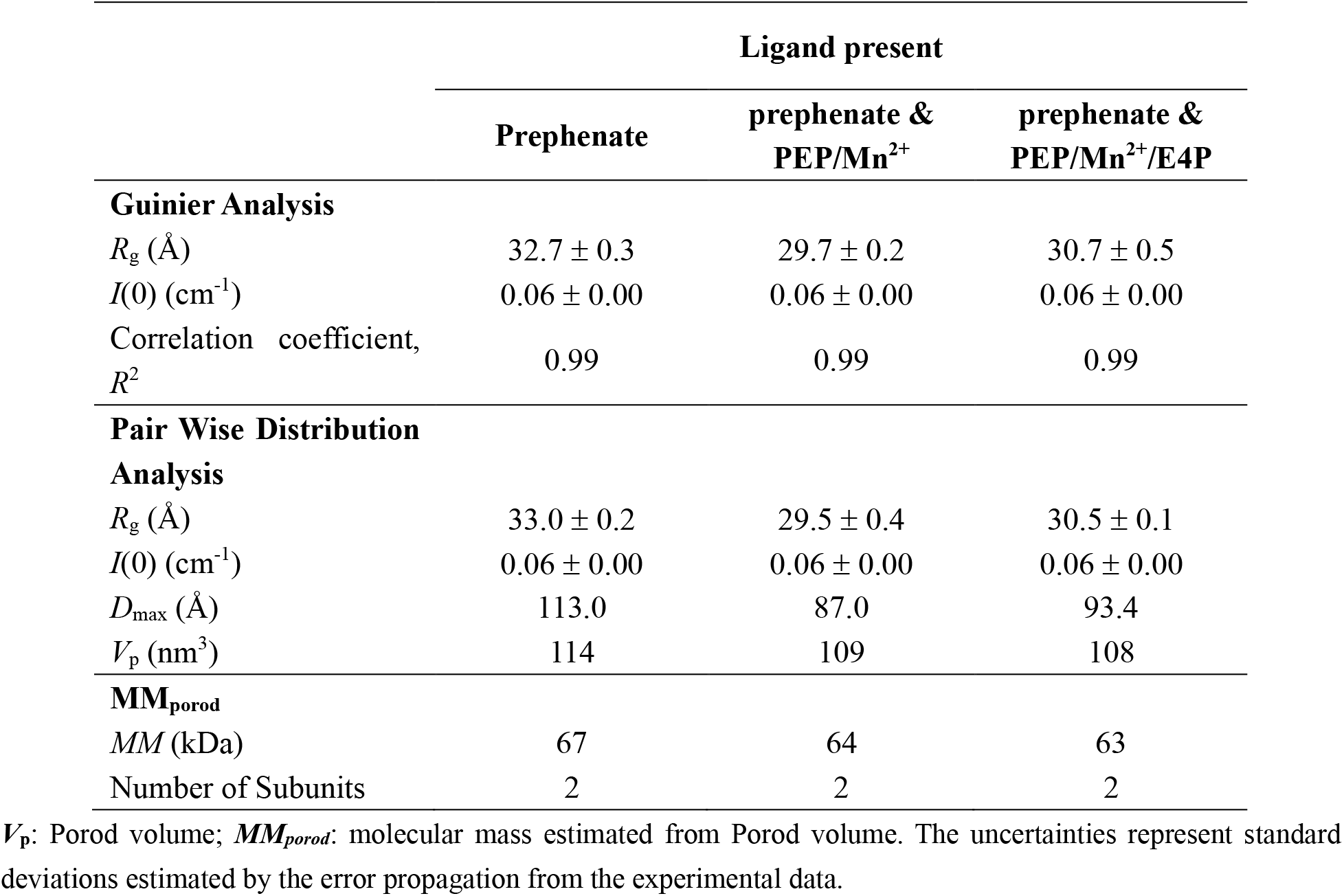
SAXS parameters of apo-*Pni*DAH7PS and *Pni*DAH7PS in the presence of PEP/Mn^2+^ or PEP/Mn^2+^/E4P, under the effect of prephenate.

|  | Ligand present |  |  |
| --- | --- | --- | --- |
|  | Prephenate | prephenate & PEP/Mn<sup>2+</sup> | prephenate & PEP/Mn<sup>2+</sup>/E4P |
| **Guinier Analysis** |  |  |  |
| $R_g$ (Å) | $32.7 \pm 0.3$ | $29.7 \pm 0.2$ | $30.7 \pm 0.5$ |
| $I(0)$ (cm<sup>-1</sup>) | $0.06 \pm 0.00$ | $0.06 \pm 0.00$ | $0.06 \pm 0.00$ |
| Correlation coefficient, $R^2$ | 0.99 | 0.99 | 0.99 |
| **Pair Wise Distribution Analysis** |  |  |  |
| $R_g$ (Å) | $33.0 \pm 0.2$ | $29.5 \pm 0.4$ | $30.5 \pm 0.1$ |
| $I(0)$ (cm<sup>-1</sup>) | $0.06 \pm 0.00$ | $0.06 \pm 0.00$ | $0.06 \pm 0.00$ |
| $D_{max}$ (Å) | 113.0 | 87.0 | 93.4 |
| $V_p$ (nm<sup>3</sup>) | 114 | 109 | 108 |
| **$MM_{porod}$** |  |  |  |
| $MM$ (kDa) | 67 | 64 | 63 |
| Number of Subunits | 2 | 2 | 2 |

$V_p$ : Porod volume; $MM_{porod}$ : molecular mass estimated from Porod volume. The uncertainties represent standard deviations estimated by the error propagation from the experimental data.

**Figure 7:**
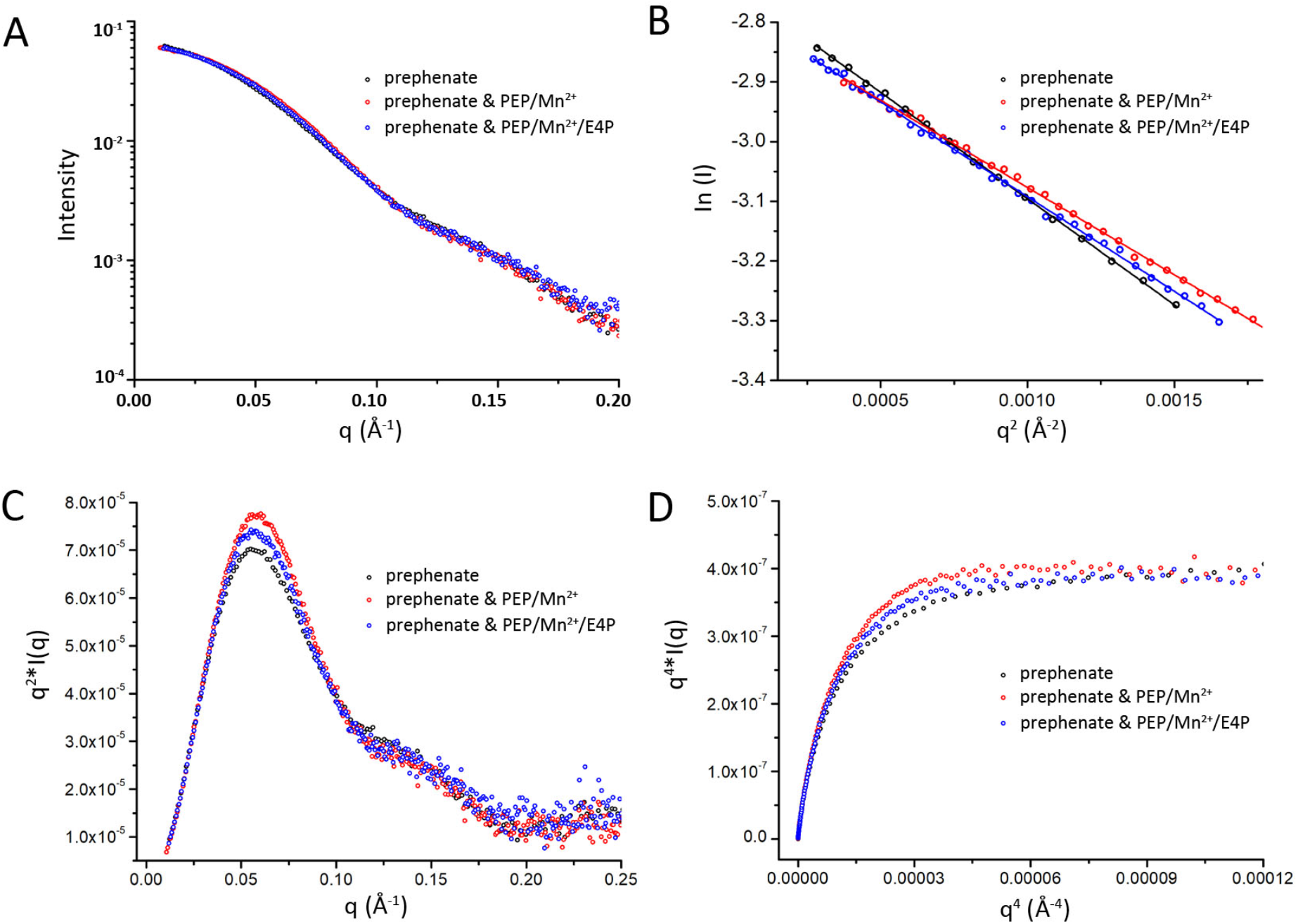
The analysis of SAXS profiles shows the structural effect of prephenate on apo-*Pni*DAH7PS (black) and *Pni*DAH7PS in the presence of PEP/Mn^2+^ (red) or PEP/Mn^2+^/E4P (Blue). **(A)** SAXS profiles (log *I*(*q*) versus *q*). **(B)** Guinier analysis (ln *I*(*q*) versus *q*^2^). **(C)** Kratky plot (*q*^2^*I(*q*) versus *q*). **(D)** Porod-Debye plot (*q*^4^\**I*(*q*) versus *q*^4^) limited to the range of the SAXS data for which the Guinier linearity was observed.

## DISCUSSION

Our previous studies of *Pni*DAH7PS uncovered an unexpected dependency between the catalytic functions of the CM and DAH7PS domains in this C-terminal CM-linked DAH7PS (26). Structural studies demonstrated that this bifunctional protein is homodimeric. In contrast to other DAH7PS enzymes, the DAH7PS domains do not directly associate and the dimer interface in *Pni*DAH7PS is only formed between the two CM domains. Therefore, the interdependency of activities between DAH7PS and CM domains in *Pni*DAH7PS can only be accounted for by direct and DAH7PS-CM interdomain interactions. In this study, we explore the conformational plasticity of this bifunctional enzyme more fully and examine the impact of substrate binding at the DAH7PS active site on the overall structure and CM catalytic function.

Scattering data demonstrates that *Pni*DAH7PS is highly flexible and does not adopt a single dominant conformation during DAH7PS catalysis. Two distinct conformational populations of the enzyme, with different DAH7PS-CM interactions, were identified, specific to the presence of either PEP/Mn^2+^ or PEP/Mn^2+^/E4P. The most compact structure of *Pni*DAH7PS was observed when PEP/Mn^2+^/E4P were present, namely when the DAH7PS-catalyzed reaction is ongoing. Intriguingly under these conditions, CM activity was substantially enhanced (Figure 2A). Therefore, we propose that the catalytic cycle of DAH7PS is intimately linked to a series of variations in DAH7PS-CM interaction, in which catalytic steps and interdomain rearrangements occur in concert to improve and fine-tune DAH7PS and CM enzymatic functions (Figure 8).

**Figure 8:**
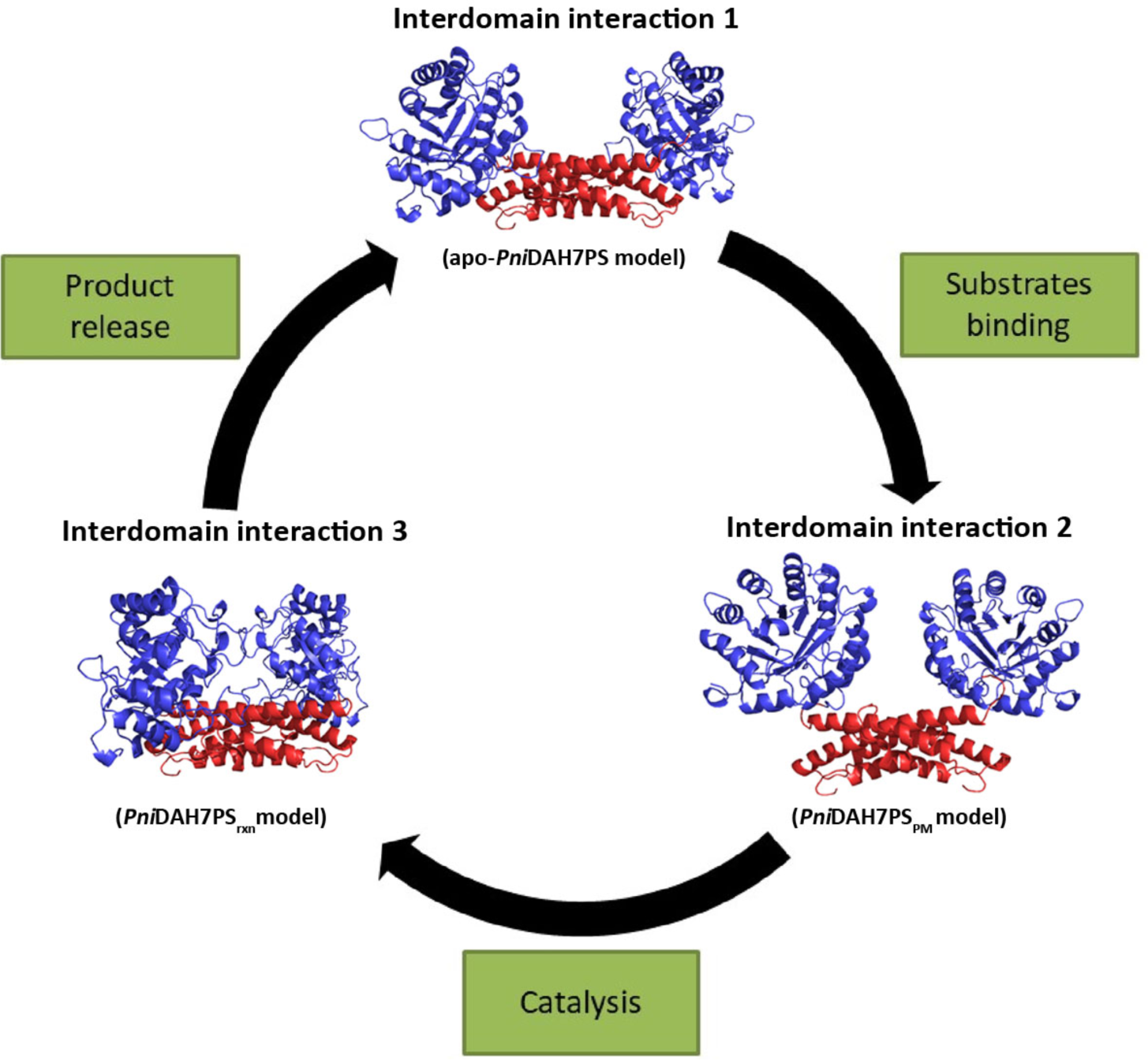
Schematic showing the hypothesized cycle of interdomain interaction of *Pni*DAH7PS throughout the DAH7PS-catalyzed reaction.

Based on sequence alignments and rigid body models for *Pni*DAH7PS (Figure 9), an extended loop within CM domain, Loop_C320-F333_, which is present in this fused DAH7PS-CM yet not in other CM types, was identified as a potential structural component that may facilitate communication to the DAH7PS domain. The functional importance of this loop was confirmed by truncation of this loop, which disrupted both the DAH7PS catalytic activity and the reaction-correlated conformational changes. Considering the complex allosteric behaviors associated with the substantial conformational changes displayed by *Pni*DAH7PS, the flexible properties of the long loop may be advantageous in assisting structural and functional variations (5).

**Figure 9:**
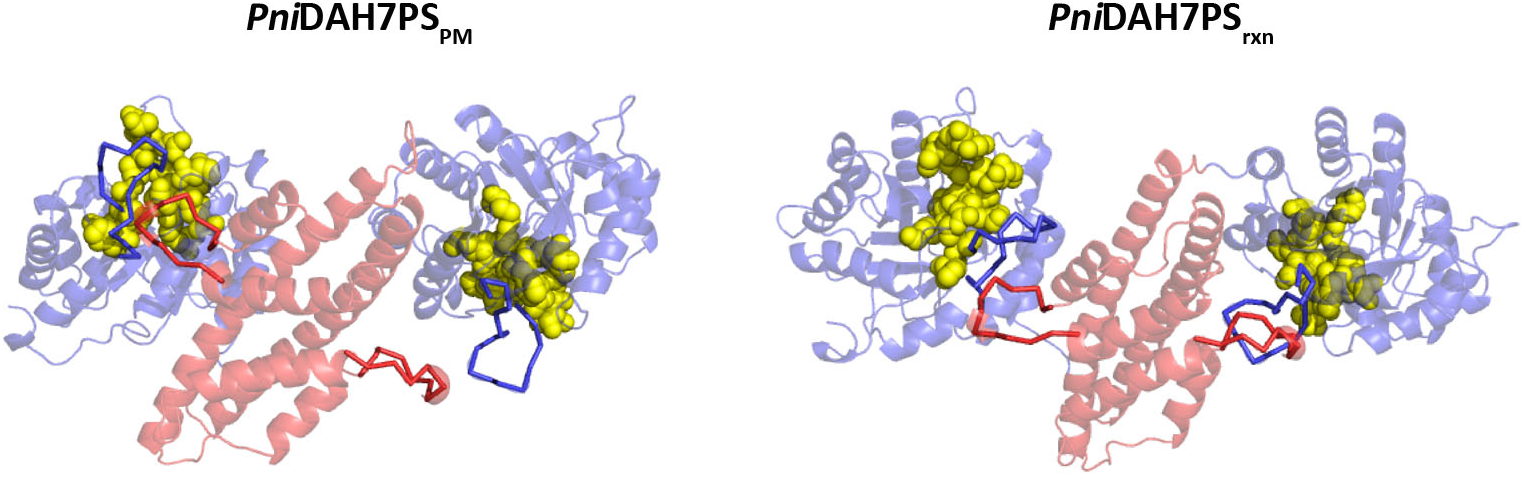
Rigid body models of *Pni*DAH7PS show the interaction between the DAH7PS (blue cartoon) and CM (red cartoon) domains respectively in the presence of PEP/Mn^2+^ (Left) and PEP/Mn^2+^/E4P (Right). The red and blue ribbon diagrams respectively indicate Loop_C320-F333_ and loop β6-α6 of DAH7PS domain that is involved in the formation of the DAH7PS active site. The yellow spheres represent the active site of DAH7PS.

Allosteric control of DAH7PS is an important strategy for metabolic regulation. Prephenate binding at the CM domain allosterically inhibits the DAH7PS activity, and the compact structure of the *Pn*iDAH7PS dimer in the presence of prephenate has been previously reported (26). Here, we observe that prephenate binding restricts the structural flexibilities of *Pni*DAH7PS associated with binding of the DAH7PS substrates. Such restrictions could attenuate the dynamic DAH7PS-CM interactions and likely disrupt the DAH7PS catalytic cycle.

Assembly of multiple enzymes in metabolic pathways is commonly found to enhance overall catalytic efficiency via co-localizing or directly channeling multiple active sites that catalyze consecutive chemical reactions (33-35). In addition, in some bi-enzyme assemblies, one functional domain/unit can play a role as the regulatory element, delivering an allosteric effect derived from its ligand binding to the other enzymatic moiety (11,20). In contrast, the bifunctional system of *Pni*DAH7PS joins together two non-consecutive enzymatic moieties; a fusion that does not serve for purpose of substrate sequestration or channeling. Furthermore, *Pni*DAH7PS displays a sophisticated interconnection between its DAH7PS and CM domains which distinguishes it from other well-studied bifunctional enzymatic systems. Specifically, in addition to the allosteric inhibition of DAH7PS by prephenate, the upregulation of CM activity in the presence of PEP/Mn^2+^/E4P delivers allosteric activation of CM mediated by the DAH7PS domain, revealing reciprocal allostery along with catalytic dependency.

This reciprocal allostery and functional interdependency of *Pni*DAH7PS provides complex feedback and feedforward response (Figure 10). Moreover, the cooperation between the feedforward activation of CM and the feedback inhibition on DAH7PS activity by prephenate allows for bidirectional regulation between the initiation and a branch point of the shikimate pathway, which enables *Pni*DAH7PS to more precisely sense both the input signals of carbon resources (PEP and E4P) and tune the output of the crucial intermediate of the shikimate pathway. Whether such complex bidirectional regulation embedded in a single multienzyme assembly is present in other biosynthetic pathways remain to be explored.

**Figure 10:**
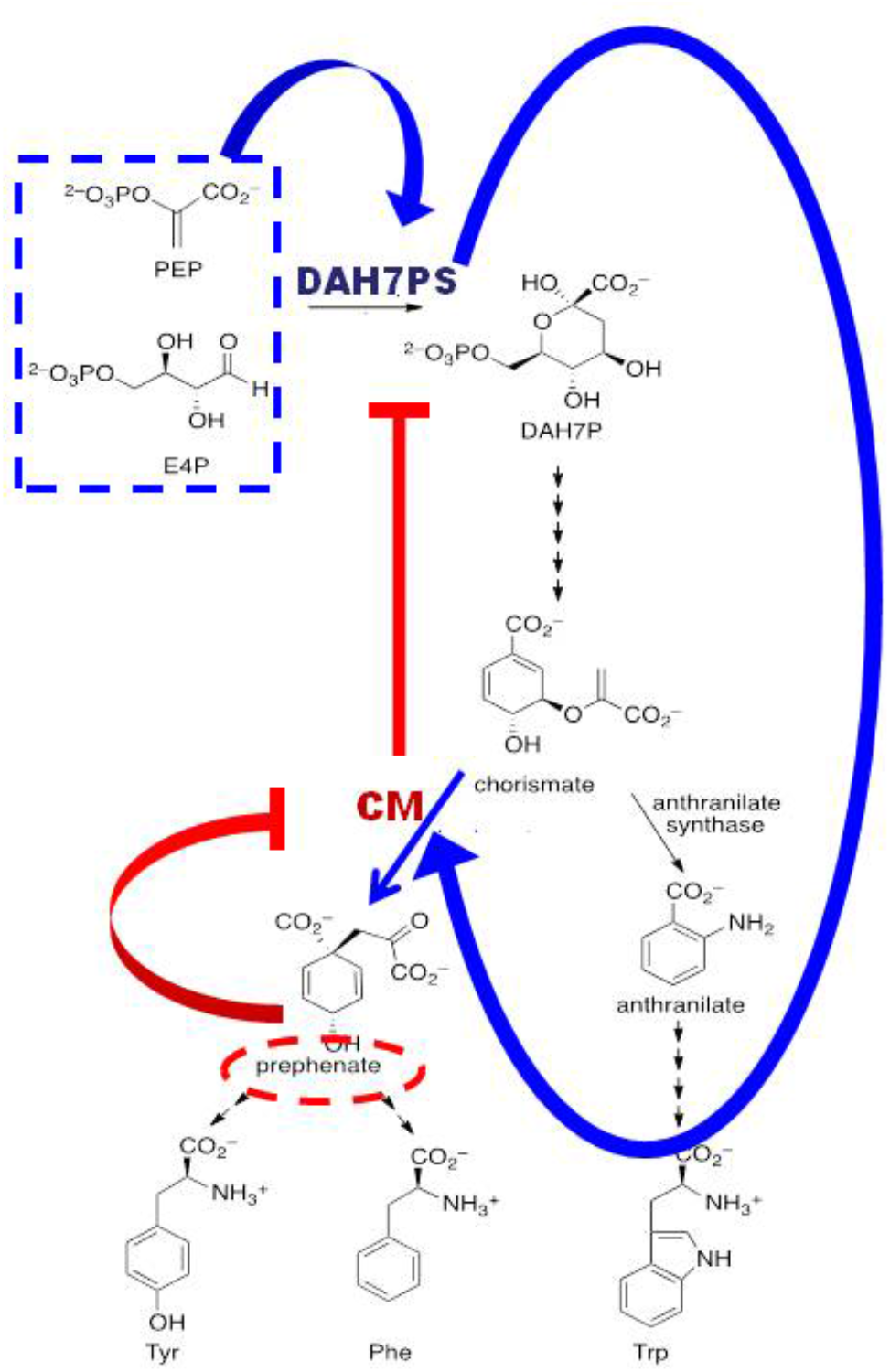
Schematic representation of the proposed bidirectional regulation mediated by *Pni*DAH7PS in the shikimate pathway. The blue point arrows represent the DAH7PS-catalyzed reaction (highlighted by blue dash rectangle) induced feedforward activation of CM activity, promoting the conversion of chorismate to prephenate; the red blunt arrows illustrate the feedback inhibition of the DAH7PS activity by prephenate (highlighted by red dash oval) via the CM domain.

## EXPERIMENTAL PROCEDURES

### Bacterial strains, plasmids, media, and growth conditions

The detailed methods for cloning, expressing and purifying the wild-type *Pni*DAH7PS have been described previously (26). Briefly, the *Pni*DAH7PS gene was cloned into pET28a vector and expressed in *E. coli* BL21 (DE3) pBB540/pBB542 cells using IPTG induction. Following sonication cell lysis, the overexpressed protein was purified sequentially using HiTrap TALON crude column (GE Healthcare) and Superdex S200 26/60 column (GE Healthcare). The purified protein was concentrated and stored at -80 °C.

### The generation of PniDAH7PS^***Δ5AA***^ ***truncated variant***

The removal of the C327-T331 segment from the CM domain of *Pni*DAH7PS was performed using a QuikChange® II Site-Directed Mutagenesis Kit (Stratagene). The pET28a-*Pni*DAH7PS plasmid was used as the template, and the deletion was generated using the primers as follows, Forward primer: 5’AAAATGTGCACGTCAGGCAAATGCAAAATTTGCAGCACGTATTCTGGAAA 3’

Reverse primer: 5’TTTCCAGAATACGTGCTGCAAATTTTGCATTTGCCTGACGTGCACATTTT 3’

PCR products were digested by *Dpn*I at 37 °C to remove the parental plasmids before they were transformed into TOP10 cells and purified. After being sequenced, plasmids with the correct deletion were transformed into *E. coli* BL21(DE3) cells. Protein expression and purification were carried out using the same methods for obtaining the purified wild-type *Pni*DAH7PS.

### Kinetic characterizations

A modified continuous assay for the spectrophotometric analysis of either DAH7PS or CM enzymatic activity (36) was employed to determine the kinetic parameters of the wild-type enzyme and *Pni*DAH7PS^Δ5AA^. Enzymatic reactions were carried out in I mL quartz cuvettes with 1 cm path length at 35 °C and the reaction rates were monitored by the disappearance of the substrate PEP at 232 nm or chorismate at 274 nm for the DAH7PS and CM activities respectively.

#### Determination of the allosteric regulation of CM activity

To detect if there would be any detectable change at 274 nm due to the additions of the components of DAH7PS reaction rather than CM-catalyzed reaction, the absorption of 500 µM PEP, 500 µM E4P, 100 µM MnSO_4_ or 500 µM DAH7P at this wavelength was checked in either the presence or absence of chorismate or *Pni*DAH7PS. The results did not exhibit any significant change in absorption. To further confirm this, a DAH7PS-catalyzed reaction was initiated and the absorbance at 274 nm was monitored, and no significant absorption was observed over time.

The determination of the effects of Mn^2+^ and the substrates of DAH7PS-catalyzed reaction on CM activity were carried out with 7.3 × 10^−2^ μM *Pni*DAH7PS using 100 µM chorismate, respectively, in the presence of 100 μM MnSO_4_, 500 μM PEP, 500 μM E4P and 500 μM DAH7P alone, or combinations of MnSO_4_ (100 μM)/PEP (500 μM), MnSO_4_ (100 μM)/E4P (500 μM) and PEP (500 μM)/E4P (500 μM). As well, the allosteric regulation of CM activity was detected as the DAH7PS-catalyzed reaction is proceeding. In this assay a DAH7PS catalyzed reaction, containing 500 μM PEP, 100 μM Mn^2+^ and 500 μM E4P, was started by either 7.3 × 10^−2^ μM *Pni*DAH7PS or *Gsp*DAH7PS. Following 5 sec incubation of the DAH7PS reaction, the CM reaction was initiated by the addition of 100 μM chorismate to determine the CM activity. 5 sec incubation makes sure that the DAH7PS-catalyzed reaction is in the linear steady-state phase (Figure S).

#### Determination of the kinetic characteristics of PniDAH7PS^Δ5AA^

The assays for determining the kinetic parameters of the DAH7PS activity for *Pni*DAH7PS^Δ5AA^ contained 100 μM MnSO_4_, 0.26 μM enzyme in 50 mM BTP buffer (pH 7.4). All reactions were initiated by the addition of E4P. Because of the high apparent *K*_M_ for E4P of *Pni*DAH7PS^Δ5AA^ and the dimerization of E4P at high concentrations (37), the equation of Alberty was employed to calculate the kinetic parameters. In this assay, the rate of consumption of PEP was measured at different fixed concentrations of PEP (167, 267 and 533 μM respectively) as E4P concentration was varied (63 – 221 μM). The data was fitted to the equation of Alberty (38):

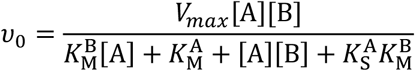

Where *V*_max_ is the maximum possible reaction rate *υ*_0_; [A] is the concentration of the variable substrate (E4P), and [B] is the fixed concentration of the other substrate (PEP) for each Michaelis-Menten kinetic measurement; *K*_M_^B^ (*K*_M_^A^) is the concentration of substrate B (A) which gives ½*V*_max_ when substrate A (B) is saturating; *K*_S_^A^ is the disassociation constant of substrate A.

The reactions for determining the kinetic characteristics for the CM domain contained 9.4 × 10^−2^ μM enzyme in 50 mM BTP buffer (pH 7.4) and initiated by the addition of chorismate with a series of concentrations (22 – 441 μM).

#### Determination of the inhibitory effect of prephenate on the wild-type PniDAH7PS

100 μM MnSO_4_, 0.26 μM enzyme in 50 mM BTP buffer (pH 7.4) were contained in the assays for determining the kinetic parameters of the DAH7PS activity of *Pni*DAH7PS. Additionally, all reactions were pre-incubated with prephenate at a saturating concentration of 100 μM before being initiated by the addition of E4P. In the presence of prephenate, the apparent *K*_M_ for E4P of *Pni*DAH7PS showed a high value, thus the equation of Alberty was adopted as well to calculate the kinetic parameters for PEP and E4P. The reaction rate was measured at different fixed concentrations of PEP (167, 267and 533 μM respectively) as E4P concentration was varied (63 – 221 μM).

### SAXS

SAXS measurements were carried out using the Australian Synchrotron (39,40) SAXS/WAXS beamline equipped with a Pilatus detector (1M, 170 × 170 mm, effective pixel size, 172 × 172 mm). The applied wavelength of X-ray was 1.0332 Å, and a sample detector, at a distance of 1.6 m from the sample, recorded scattered X-rays intensities covering the *q* range of 0.0015-3.0 Å^-1^. Either wild-type *Pni*DAH7PS or *Pni*DAH7PS^Δ5AA^ was eluted from a SEC column (Superdex 200 5/150) using SEC buffer (10 mM BTP, pH 7.4, 150 mM NaCl and 3% (v/v) glycerol) and the buffers containing various additives of interest as listed in Table S1. The eluted sample passed into a 1.5 mm thin-walled glass capillary where the sample was detected using X-rays at 25 °C at 2 s intervals. The raw scattering data was processed using Scatterbrain, developed at the Australian Synchrotron, for data reduction and buffer subtraction. Processed data were plotted as *I*(*q*) vs. *q* (*q* = (4πsinθ)/ λ, 2θ is the scattering angle), and analyzed using Primus (41).

All data sets for structural analyzes were recorded with ∼ 400 data points over the range 0.005 ≤ *q* ≤ 0.35 Å^-1^, and a single symmetric scattering peak over ∼ 120 frames was observed in each data set. To confirm conformational homogeneity, three different groups of data respectively covering the maximum scattering intensity and either side of the scattering maximum were summed and compared. The same scattering profile was observed across each scattering peak. The radius of gyration (*R*_g_) of the protein particle was evaluated from Guinier plots (ln*I*(*q*) vs. *q*^2^) of scattering data. In addition, the linear Guinier plot in the *q* range of *q* ≥ 1.3/*R*_g_ was a crucial indicator of the monodispersity of samples. The Porod volume (*V*_p_) and the maximum distance (*D*_max_) of the protein particle were evaluated respectively from the Porod plot and the pair-distance distribution (*P*(*r*)) function obtained via Fourier transformation. *MM* _*porod*_ was obtained by dividing the Porod volumes by 1.7.

### Homology and Rigid body Modelling

Homology modelling of monomeric *Pni*DAH7PS^D^ and dimeric *Pni*DAH7PS^CM^ was performed by Modeller using *Pfu*DAH7PS (42) and *Pfu*CM (43) as template structures, respectively (44). Then, the two resultant homology models were connected together as a full length monomeric unit (*Pni*DAH7PS-CM) containing an undefined linker using the software PRE_BUNCH (45). Subsequently, two *Pni*DAH7PS-CM monomeric models were docked into the SAXS *ab initio* envelope of apo-*Pni*DAH7PS, *Pni*DAH7PS_PM_ and *Pni*DAH7PS_rxn_ to generate a rigid body model using BUNCH, constrained by two-fold symmetry and distance information of the contacting residues obtained from the homology model of the *Pni*DAH7PS^CM^ dimer (Table S2).

## DATA AVAILABILITY

All data are given in the main manuscript or supporting information.

## ACKNOWLEDGEMENTS

Data were collected at the SAXS beamlines of the Australian Synchrotron (39,40) with access provided by the New Zealand Synchrotron Group.

Dr. Wanting Jiao (Ferrier Research Institute, Victoria University of Wellington) reviewed this manuscript and provided valuable comments.

## AUTHOR CONTRIBUTIONS

YB carried out the experiments and bioinformatics analysis, EJP and YB analyzed the experimental data, EJP conceived and coordinated the study, YB and EJP wrote the paper. All authors reviewed the results and approved the final version of the manuscript.

## FOOTNOTES

### Funding: Marsden Fund VUW1426

#### The abbreviations used are

DAH7PS, 3-deoxy-d-*arabino*-heptulosonate 7-phosphate synthase; CM, chorismate mutase; *Pni*DAH7PS, the C-terminal CM-linked DAH7PS from *Prevotella nigrescens* DAH7PS; *Pni*DAH7PS^D^, the DAH7PS domain of *Pni*DAH7PS; *Pni*DAH7PS^CM^, the CM domain of *Pni*DAH7PS; *Pgi*DAH7PS, the C-terminal CM-linked DAH7PS from *Porphyromonas gingivalis*; *Pgi*DAH7PS^CM^, the CM domain of *Pgi*DAH7PS; *Gsp*DAH7PS, the N-terminal CM-linked DAH7PS from *Geobacillus sp*.; *Mtu*DAH7PS-CM, DAH7PS-CM complex from *Mycobacterium tuberculosis*; *Pfu*CM, the monofunctional CM from *Pyrococcus furiosus*; *Ape*CM, the monofunctional CM from *Aeropyrum pernix*; E4P, erythrose 4-phosphate; PEP, phosphoenolpyruvate; SEC, size-exclusion chromatography; SAXS, Small Angle X-ray Scattering; *Pni*DAH7PS_PM_, the PEP/Mn^2+^ bound *Pni*DAH7PS; *Pni*DAH7PS_rxn_, the *Pni*DAH7PS in the presence of PEP/Mn^2+^/E4P; *Pni*DAH7PS^Δ5AA^, a truncated *Pni*DAH7PS variant with removal of C327-T331 residues; 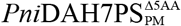, the PEP/Mn^2+^ bound *Pni*DAH7PS^Δ5AA^ ; 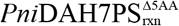, the *Pni*DAH7PS^Δ5AA^ in the presence of PEP/Mn^2+^/E4P

